# Heterogeneous chromatin mobility derived from chromatin states is a determinant of genome organisation in *S. cerevisiae*

**DOI:** 10.1101/106344

**Authors:** Sven A. Sewitz, Zahra Fahmi, Latifa Aljebali, Jeremy Bancroft, Otávio J. B. Brustolini, Hicham Saad, Isabelle Goiffon, Csilla Várnai, Steven Wingett, Hua Wong, Biola-Maria Javierre, Stefan Schoenfelder, Simon R. Andrews, Stephen G. Oliver, Peter Fraser, Kerstin Bystricky, Karen Lipkow

## Abstract

Spatial organisation of the genome is essential for regulating gene activity, yet the mechanisms that shape this three-dimensional organisation in eukaryotes are far from understood. Here, we combine bioinformatic determination of chromatin states during normal growth and heat shock, and computational polymer modelling of genome structure, with quantitative microscopy and Hi-C to demonstrate that differential mobility of yeast chromosome segments leads to spatial self-organisation of the genome. We observe that more than forty percent of chromatin-associated proteins display a poised and heterogeneous distribution along the chromosome, creating a heteropolymer. This distribution changes upon heat shock in a concerted, state-specific manner. Simulating yeast chromosomes as heteropolymers, in which the mobility of each segment depends on its cumulative protein occupancy, results in functionally relevant structures, which match our experimental data. This thermodynamically driven self-organisation achieves spatial clustering of poised genes and mechanistically contributes to the directed relocalisation of active genes to the nuclear periphery upon heat shock.

**One Sentence Summary:** Unequal protein occupancy and chromosome segment mobility drive 3D organisation of the genome.

## Main Text

Eukaryotic genomes are highly organised in three dimensions (*1, 2*) and this spatial organisation has to be maintained in order to achieve the correct gene expression profiles (*3–6*). The 3D organisation of the genome is thus central to many aspects of cell biology and has been intensely investigated during normal growth (*7–9*), differentiation (*10–12*), cell division (*13*), senescence (*14*), and disease (*5, 6, 15*), and has been shown to arise independently of transcription (*16*). In the budding yeast *Saccharomyces cerevisiae,* target genes of most transcription factors are enriched in specific regions along the chromosome in one dimension (*17*), or in the genome in three dimensions (*18*). A central question in the field is by which mechanisms this 3D organisation is achieved.

Any mechanism that organises genome structure has to do so in a highly dynamic and crowded nucleoplasm (*19, 20*). The prevalent view is that 3D genome organisation comes about despite the known intrinsic fluctuations of the chromatin fibre. Most studies focus on stable interactions between DNA-bound proteins that connect two chromatin loci (*9, 21–27*). Here, we propose and validate a fundamentally different mechanism: The mobility of the chromatin fibre is not uniform, but heterogeneous, along its length, as a result of the unequal distribution of protein binding along the genome. This leads to thermodynamically driven self-organisation, which we observe experimentally, and which we show to have important functional implications.

## Determination and characterisation of chromatin states

In order to analyse the global effects of protein binding on spatial organisation of yeast chromosomes, and incorporate these data into a computational model, we determined chromatin states in yeast. Chromatin states (also named ‘chromatin colours’) are an important conceptual advance in the field of chromatin biology. Here, chromatin modifications (*28*), chromatin-associated proteins (*29*), or a combination thereof (*30, 31*) are functionally categorised into groups or states, giving a chromatin-centric annotation of the genome. The resulting chromatin states were shown to correspond to differences in transcriptional activity, including the developmental regulation of genes (*29–31*) and 3D genome organisation (*21, 22, 32–34*).

To determine chromatin states for the budding yeast *Saccharomyces cerevisiae,* we modified the method of (*29*) to employ yeast chromatin immunoprecipitation (ChIP) data as input (*35*). This method determines chromatin states from quantitative protein binding data alone. We used the reported genome-wide binding profile of 201 chromatin-associated proteins, measured in cells grown at 25°C, and 15 minutes after shifting the culture to 37°C (heat-shock) (*36*) (Fig. 1, Fig. S1). Five states effectively differentiated the protein binding profiles between the states (see SI for more details). The same procedure was performed independently for the 25°C and 37°C data. The genes in each state were counted and the states were numbered S1-S5 according to their decreasing coverage at 25°C (Fig. S1D, Fig. S2A).

**Fig. 1.**
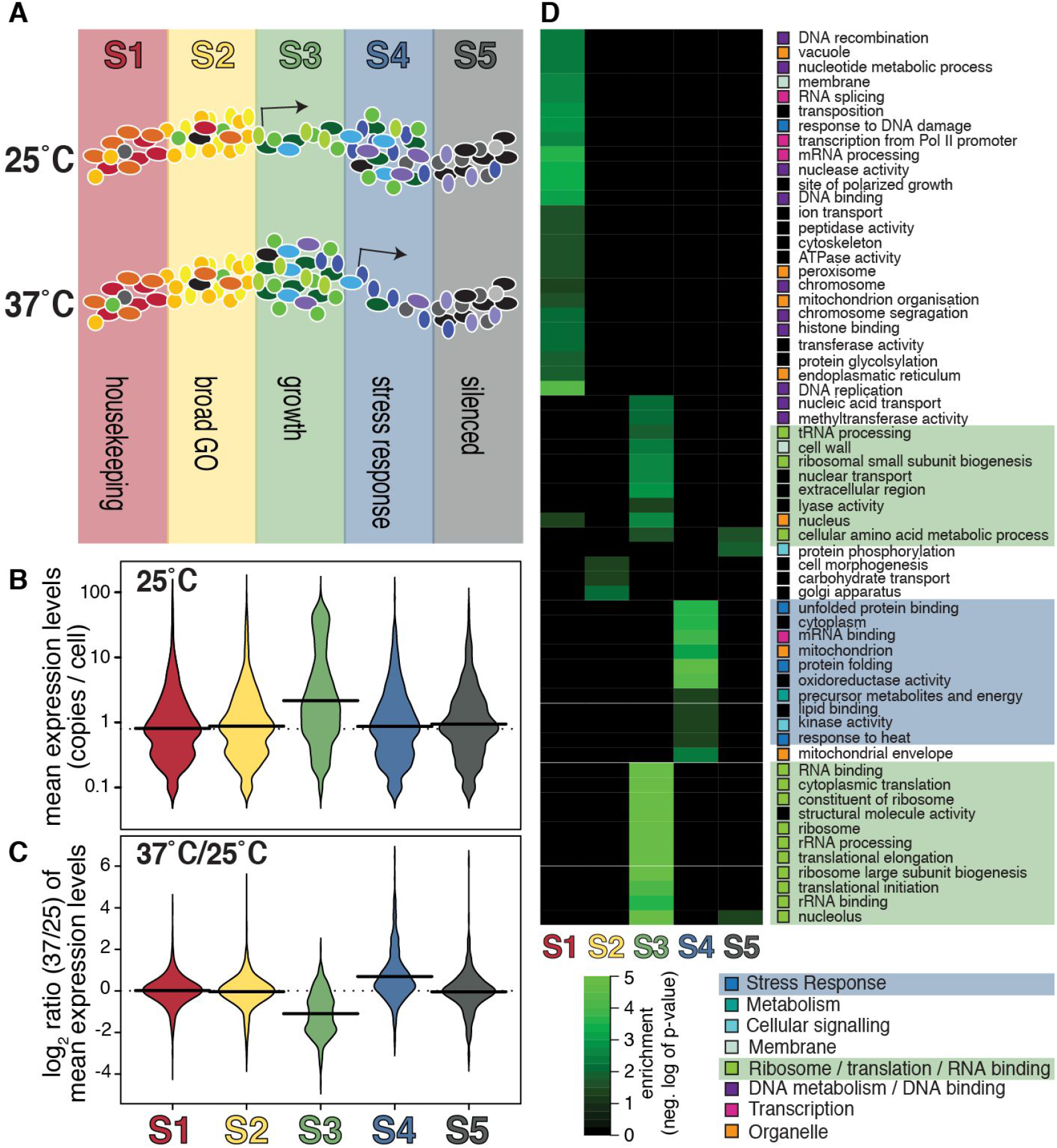
Chromatin states in *S. cerevisiae* and their characteristics. **(A)** Overview: We defined chromatin states using statistical methods, from the combination of 201 bound chromatin-associated proteins (coloured ellipses) (see Fig. S1). The states were ordered by genome coverage (highest: S1 to lowest:S5), and assigned a colour (red, yellow, green, blue, grey). At 25°C, the genes in S4 have a greater amount of proteins bound, while genes in S3 have higher expression (kinked arrow). At 37°C, this is reversed, with genes in S3 showing higher protein occupancy, and S4 genes higher expression. A high-level gene ontology analysis per state is shown. **(B)** Expression analysis of chromatin states from yeast grown at 25°C. Thick horizontal lines: mean; dashed line: total average. At 25°C, S3 (green) has a significantly higher number of highly expressed genes (p<2.2e-16); expression array data from (*41*). **(C)** After shift to 37°C, the ratio of expression (37°C/25°C) shows that genes in S3 are repressed, while those in S4 are upregulated; expression array data from (*42*). **(D)** Gene Ontology enrichment analysis of genes in each state, using Ontologizer (*43*).

## Functional analysis of chromatin states

At 25°C, S3 contains more highly expressed genes and significantly higher median expression than all other states (Fig. 1B). At 37°C, expression is reduced by 53% in S3, while expression of genes in S4 is increased by 62%. Expression of genes in S1, S2 and S5 remains largely unchanged (Fig. 1C; Table S2, column 5).

S1 harbours categories typical of housekeeping genes (*37*) (Fig. 1D). Genes in S2 show very limited GO enrichment, indicating that these genes are distributed over many functional categories. Genes assigned to S3 cover diverse processes and functions necessary for maintaining high levels of translation during rapid growth. Genes in S4 include functions such as unfolded protein binding, response to heat, and protein chaperones; all typical for heat-shock response genes. S5 is enriched for protein phosphorylation, amino acid metabolism and the nucleolus; it also harbours genes that code for proteins located at telomeres, such as the telomere-binding protein Cdc13p (*38*). These ontologies match the change in expression profile observed upon heat-shock, as genes required for translation (S3) are known to be highly expressed under favourable conditions and to be repressed when cells are stressed (*39, 40*), while heat-shock genes (S4) are expected to be upregulated at 37°C (Fig. 1A-C).

## Protein occupancy and poising

We investigated the distribution of proteins across chromatin states. For each protein, we applied a binary hidden Markov Model (HMM) to the raw ChIP data to identify occupied sites and classified them by state (see (*35*)). We calculated the ‘fraction occupied’ for all states and all proteins, at both temperatures (Fig. S3).

The three RNA polymerase II subunits present in the ChIP dataset, Rpb2p, Rpb3p and Rpb7p (Fig. 2A), display different binding patterns across chromatin states. Their fraction occupied values per state are shown in Fig. 2B. Interpretation is made easier when assessing the rank protein occupancy (Rank occupancy, Fig. 2C). At 25°C, Rpb2p and Rpb3p have the highest binding rank to S4 genes, and the lowest binding rank to S3 genes. The rank order for both these RNA polymerase subunits is identical, reflecting the close contacts they make within the RNA Pol II enzyme (Fig. 2A). After the shift to 37°C, both Rpb2p and Rpb3p relocate, and the ranking of S3 and S4 is reversed: now S3 is bound most (highest rank), and S4 the least (lowest rank) (Fig. 2C). The rank order of both Rpb2p and Rpb3p is again identical. This is in stark contrast to the rank occupancy for Rpb7p: At 25°C, the highest occupancy is in S3, and at 37°C in S4 (Fig. 2C). At both temperatures, the preferred location of Rpb7p coincides with the chromatin state which shows the highest level of expression (Fig. 1B). The Rpb2p and Rpb3p subunits, however, show a distribution that is termed ‘poised’: ready for immediate activation (*44*). This means that the highest levels of binding occurs at the state that becomes active under different temperature conditions. Note that here poised genes are defined by protein occupancy and not by histone modification (*45*).

**Fig. 2.**
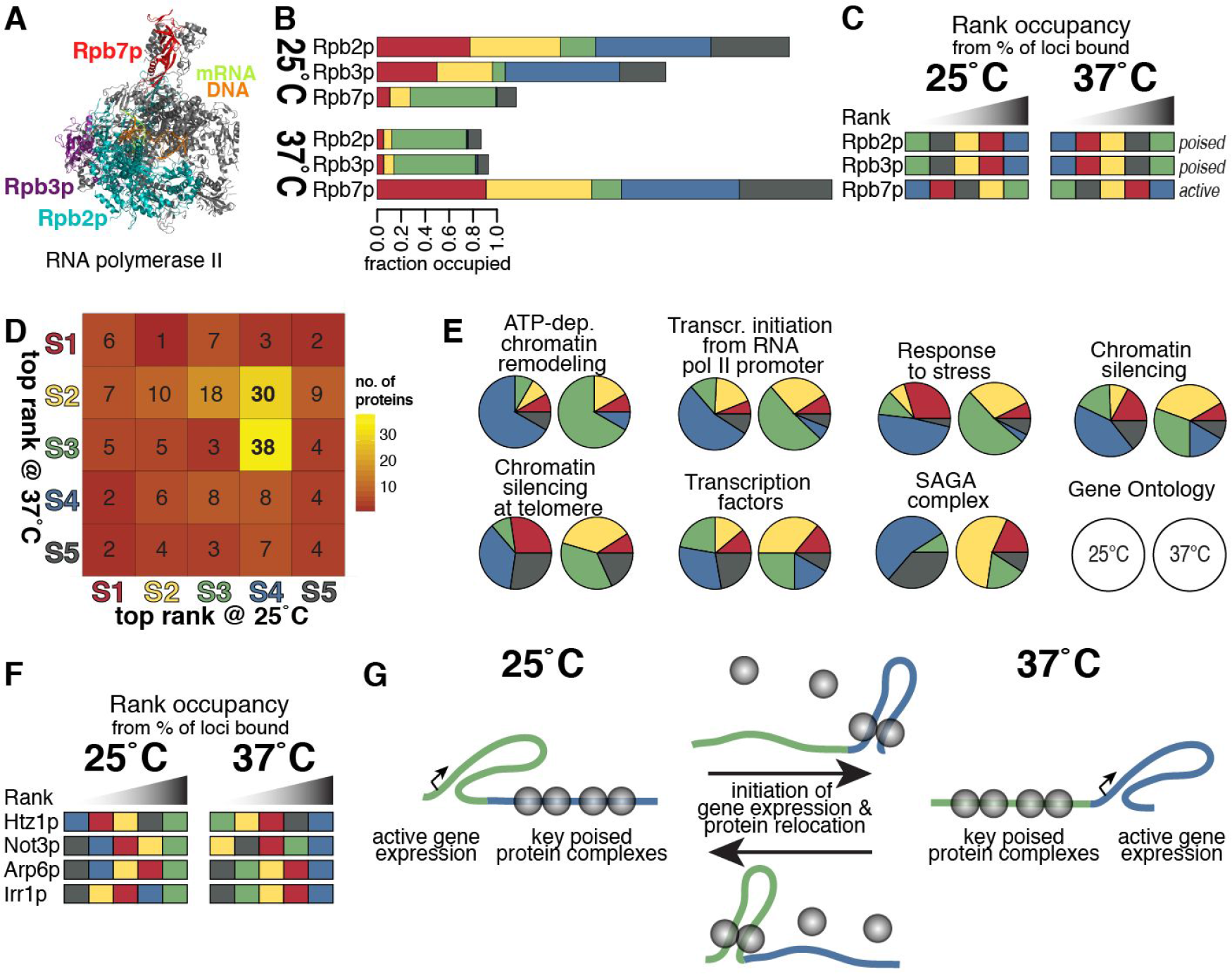
Wide-spread, consistent poising in the yeast genome. **(A)** Structure of RNA polymerase II in complex with DNA and RNA transcript. Crystal structure from PDB 3HOU (*57*). RNA Pol II is a complex of 12 proteins, Rpb1p to Rpb12p. The three proteins included in the analysed dataset (*36*) are coloured: Rpb2p (teal), Rpb3p (purple) and Rpb7p (red). Note that Rpb7p is part of a small subcomplex sitting on top of the main complex. **(B)** Stacked barplots of the fraction occupied values for the three RNA Pol II subunits, for 25°C and 37°C (from Fig. S3). **(C)** Rank occupancy plot. At 25°C, both Rpb2p and Rpb3p bind most to genes in S4, indicated by the blue square to the right, while the Rpb7 subunit is bound most to S3. At 37°C, this pattern is reversed. Rpb2p and Rpb3p are described as ‘poised’. The binding location of Rpb7p coincides with actively transcribed genes (‘active’). **(D)** Top-ranked binding state of all 201 proteins at 25°C and 37°C. **(E)** Poising across functional protein types. The pie charts show the distribution of proteins in each family across the chromatin states, taking into account only the highest rank for each protein. **(F)** Proteins that are most bound to active genes at both temperature conditions (from those with ≥20% difference in occupancy, Fig. S3). **(G)** Schematic showing how gene expression (kinked arrow) and poising changes under different growth conditions.

The difference between subunit distributions (Fig. 2A) can be explained by the structural composition of the 12-subunit RNA Pol II enzyme. Rpb2p and Rpb3p make extensive contacts to each other and are part of the core enzyme complex, while Rpb7p binds only minimally to the core complex. Rpb7 (together with Rpb4) is in a diffusible subunit, known to be present in substoichiometric concentrations (*46*), and to participate in the stress response (*47, 48*). RNA Pol II has been reported to be poised during stationary phase (*44*). Our analysis demonstrates that a large fraction of the main RNA Pol II complex is poised, both in exponential phase at favourable temperatures (25°C), and during heat stress (37°C), and that the location of the poised polymerase changes. Our results also confirm that the presence of Rpb7p is linked to actively transcribed genes (*49*).

We investigated whether other proteins also showed a poised distribution. At 25°C, 85 proteins (42%) have their highest levels of occupancy in S4. At 37°C, 38 of these poised proteins binding with the highest rank to S3 and 30 proteins to S2 (Fig. 2D). This indicates that significantly more proteins than previously described are poised, and that much of the previously described widespread movement of proteins upon heat-shock (*36*) involves a concerted migration from S4 to S3 and, to a lesser extent, to S2 (Fig. 1A, 2G).

Grouping the proteins according to molecular function shows that not all are poised to the same degree (Fig. 2E). We here define a protein as being poised if its highest rank is S4 at 25°C and S3 at 37°C. ATP-dependent chromatin remodellers show the highest level of poising (66.7%, Fig. 2E), followed by the protein components responsible for transcription initiation from an RNA Pol II promoter (54.5% at 25°C, 51.2% at 37°C). Transcription factors show the lowest levels of poising (19.4%). Hence, we conclude that chromatin remodellers hold the gene in an activation-ready state, with transcriptions factors acting to trigger gene expression and only binding at the time when the gene product is required.

Of the 201 proteins analysed, the rank occupancy of only five proteins that differ by more than 20% between any two states, correlated positively with transcriptional activity at both temperatures: Htz1p, Not3p, Arp6p, Irr1p and Rpb7p (Fig. 2C,F). All these proteins have well established roles at sites of active transcription (*50–56*).

## Polymer model of mobile chromosome organisation

The significant differences in protein occupancy between segments of chromatin, as determined by chromatin states, justify modelling chromatin as a heteropolymer. We postulated that the heteropolymeric nature of chromatin would affect the local mobility of each segment. In this context, the term mobility describes the average linear displacement of a genome segment at each time step, i.e. the instantaneous velocity of its Brownian motion. We further postulate that this heterogeneity of segment mobilities affects the structure of the whole genome.

In order to investigate this new concept, we carried out polymer simulations, in which we assigned different mobilities to segments according to their chromatin state. To this end, we modified a previously developed and validated three-dimensional computational polymer model of the interphase yeast genome (*58*). In this model, each chromosome is a polymer of cylindrical segments, connected by ball joints, attached to the spindle pole body at the centromere, and constrained by the nuclear membrane (see further details in (*58*) and in the SI). We reduced the length of each segment to 2 kb, which is the approximate average length of a gene and its flanking sequences in *S. cerevisiae (59)*, and assigned the corresponding chromatin state to each segment of the computational model (Fig. S4A, Fig. 3, Movie S1).

**Fig. 3.**
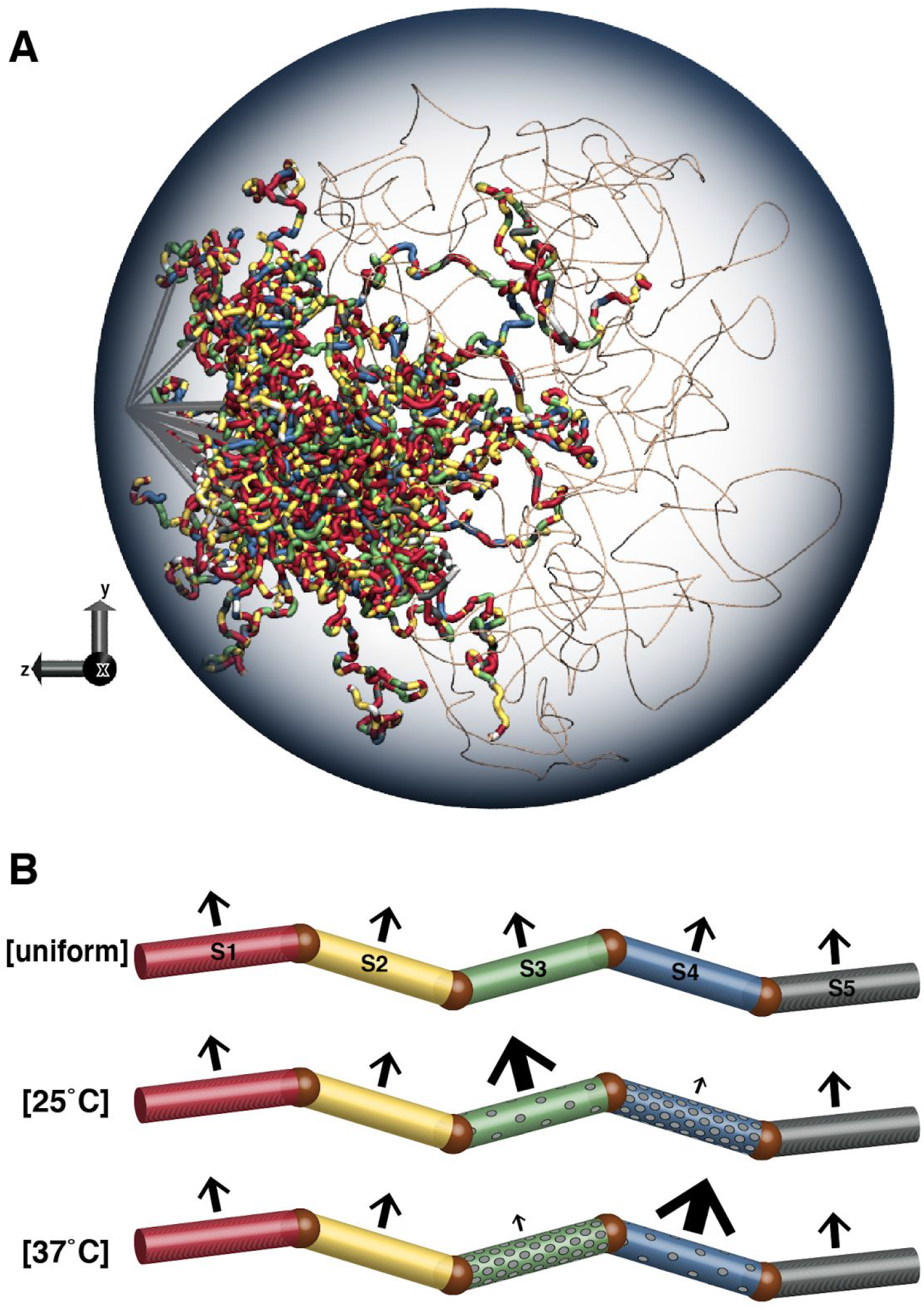
Computational whole genome model. **(A)** Snapshot of the stochastic model, with 2 kb cylindrical segments and state mapping. Centromeres are attached to short microtubules (straight grey tubes) at the spindle pole body. The segments shown as thin beige lines represent the rDNA repeats and form the nucleolus (thickness not drawn to scale). For state assignment and measurements of all segments, see Fig. S4A and Table S3. Following the convention of the literature on *S. cerevisiae* genome structures, this plot and subsequent figures are rotated such that the centromeres are shown on the left. **(B)** Each genome segment is assigned the appropriate chromatin state. To simulate different protein occupancies, the compound Langevin force (*F_LC_*) that is applied to each segment is varied according to chromatin state. Representing the 25°C situation, *F_LC_* is reduced for all S4 segments (small arrow) and increased for all S3 segments (large arrow). For clarity, all arrows are pointing upwards; in the simulation, *F_LC_* vectors are calculated stochastically and independently for each segment and each dimension at every time step.

Proteins can affect chromatin mobility through a number of different physical properties by e.g. changing the diameter, charge, viscosity, and transient interactions of the chromatin fiber. For a measure that encompasses all these influences, we chose to directly modify the force that drives segment mobility. In this model, as in most others, the mobility of genome segments in three dimensions is governed by Langevin dynamics, which is composed of the random force (Brownian dynamics), and the viscous drag (*58, 60*). We call this the compound Langevin force, *F_LC_*. The *F_LC_* is calculated independently for each segment at every time step. To implement differential mobility of chromatin states, we multiplied the thus calculated force by a state-specific factor (Fig. 3B).

We investigated the relationship between protein binding and the compound Langevin force in short test simulations (Fig. S5): Increasing the mass and radius (i.e. the protein occupancy) of a segment has the same qualitative effect on a segment’s displacement per time step as decreasing the *F_LC_*, and *vice versa* (compare matching columns in Fig. S5B with C). Thus, changing the *F_LC_* has the desired effect of changing the mobility of genome segments in line with protein occupancy.

As shown by our state analysis of protein binding data, the largest change in protein occupancy occurred from S4 at 25°C to S3 at 37°C (Fig. 2D, summarised in Fig. 2G). We therefore limited our analysis to reciprocally changing the forces applied to segments of these two states (Fig. 3B). The uniform (homopolymer) model, in which no factor is applied to any calculated *F_LC_*, served as control. For analysis, we let the simulations reach steady state and then collected data from non-correlated time steps (Fig. S6).

## Experimental validation of the model: Microscopy

To experimentally validate our computational model, we first used live cell fluorescence imaging. We created a series of yeast strains in which two sites at genomic distances varying between 27 kb and 495 kb of the left arm of chromosome XIV are tagged by the lac and tet fluorescent operator systems (Fig. 4A(a,b), Table S4). Fluorescent signals are generated by GFP- and mRFP- fluorescently labelled lac and tet repressor proteins bound to their respective operator repeat sequences. Images were acquired in 3D (stacks of 21 images at z = 200 nm) and distances between the tagged sites determined using an automated ImageJ-based algorithm (*61*) (Fig. S7). Measured median distances ranged from 422 nm to 765 nm for loci separated by up to 220 kb. At greater genomic distances, the measured median did not increase further (Fig. 4A(d)). We compared these distance distributions with distributions obtained from simulations (Fig. 4A(e-g)). The uniform model predicted median distances between loci separated by 27 to 495 kb ranging from 125 nm to 720 nm (Fig. 4A(e)). For small genomic distances (27-79 kb), the simulated values were significantly smaller than the measured physical 3D distances due to experimental noise (Fig. 4A(c), (*62*)). In contrast to the *in vivo* results, however, a uniform polymer model produces distances which increase monotonically with genomic separation without reaching a clear plateau (Fig. 4A(e)). Hence, we simulated distance distributions using a series of heteropolymeric models, with different sets of state-specific factors to change the *F_LC_* exerted on segments. The [25°C] models in which the forces applied to the S3 and S4 segments are reciprocally changed 5-fold (Fig. 4A(f,g)) generate distance distributions that are statistically the closest to the experimental data (non-nested Vuong closeness test (*63*), see details in SI, Table S7). Strikingly, the models with 5-fold changes are also the only ones that do not show a strict monotonic rise, with the 220 kb and 319 kb positions showing near identical 3D distances. Simulations based on 2-fold or 10-fold change in *F_LC_* on any of the segments (Fig. S8A) show a statistically lower agreement with the *in vivo* data (Table S7). This analysis shows that the heteropolymeric model simulating the [25°C] conditions is a better fit to the experimental data than both the homopolymeric model and the model simulating [37°C] conditions.

**Fig. 4.**
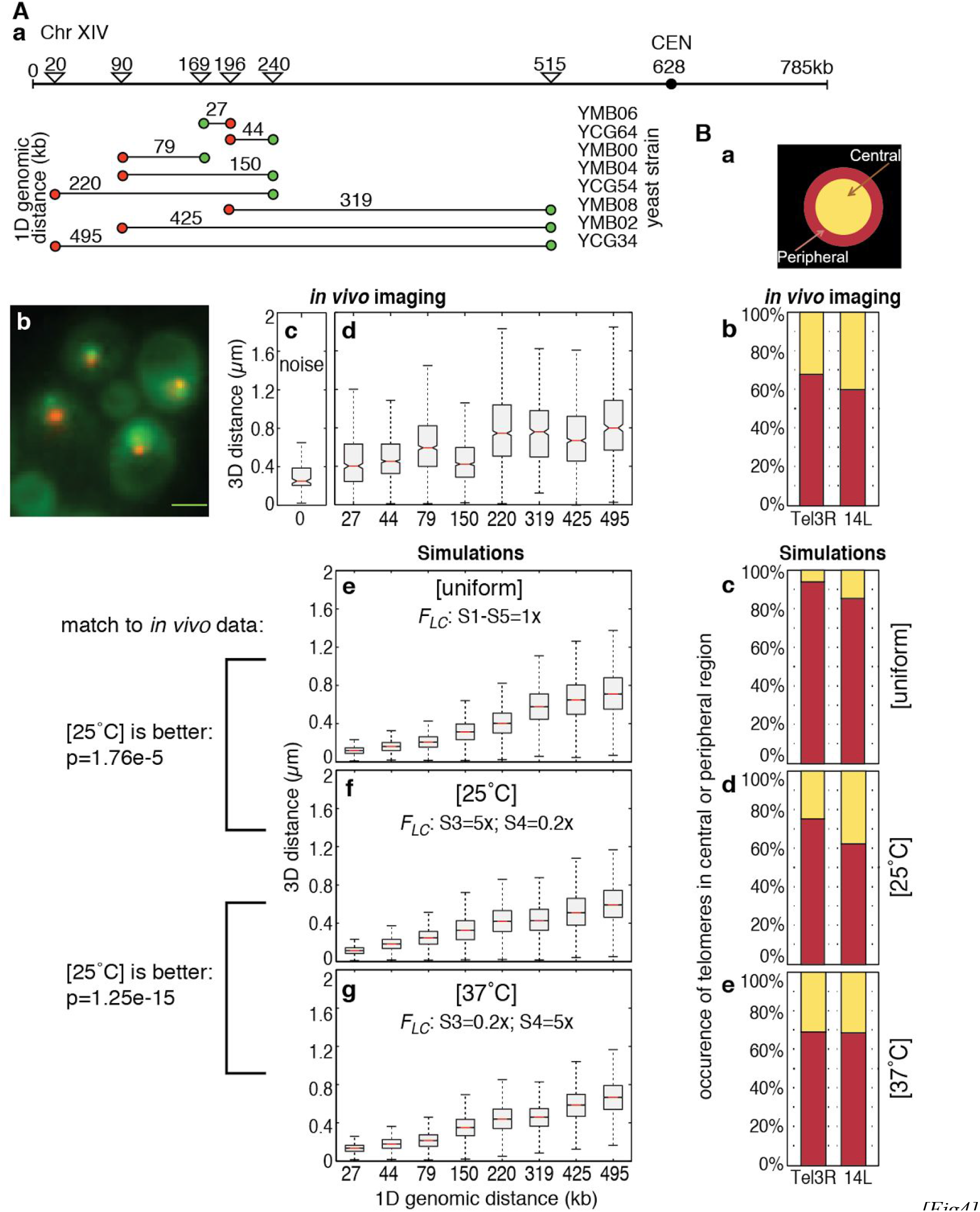
Validation of the model by microscopy. Position of labelled gene loci in confocal microscopy and in simulations. **(A)** Genomic loci at different distances from each other, and at different distances from the chromosome end, were labeled with the fluorescent repressor operator system (FROS) on the left arm of Chromosome XIV. **a)** Positions of the insertions in kb, recognised by mRFP- and GFP-labelled fusion proteins. The distances between the genomic positions of the operator array insertions (27 kb to 495 kb) are shown above the lines connecting the red and green labels (see also Table S4). **b)** Representative wide-field image, maximum projection of strain YCG54 (“220 kb”). Scale bar: 2 μm. **c)** Distribution of measured 3D distances in strain YPT237 (*62*), in which a single array is recognised by both mRFP- and GFP-fusion proteins: Control for experimental noise. **d)** Distribution of distances per yeast strain (n=498 to n=1233). **e)** Distribution of distances between mapped loci in uniform (homopolymer) simulations. **f,g)** Distribution of distances in heteropolymer simulations. Models were compared using Vuong tests, see Table S7. **(B)** Telomere positions. **a)** The nuclei were divided into a central and peripheral zone of equal area. **b)** Position of labelled telomeres in live cells; n=80 (Tel3R); n=74 (Tel14L) (modified from (*20*)). **c-e)** Telomere positions in the simulated nuclei; n=30,000. The *in vivo* data is significantly different from the uniform simulation data (binomial test: p=3.9e-12 (Tel3R) and p=2.2e-7 (Tel14L)). It is not significantly different from the heterogeneous simulation data (p>0.15), and the heterogeneous simulation data only is within the confidence interval of the *in vivo* data (Table S8).

We next assessed whether our model could predict specific features of nuclear organisation. As it has been shown that budding yeast telomeres are preferentially, but not systematically, located near the nuclear membrane (*19, 64–66*), we recorded the frequency of peripheral localisation of two different telomeres. Applying the uniform model, positions of the right telomere of Chr III (Tel3R) and the left telomere of Chr XIV (Tel14L) were located in the most peripheral zone in 94.0% and 85.3% of the sampled time-steps, respectively (Fig. 4B(c), Table S8). These numbers are significantly different from our previously reported measurements of live cells (*20*), where only 68.8% and 60.8% of these telomeres were found in the periphery (Fig. 4B(b), Table S8). In contrast, in simulations based on heteropolymeric models with 5x reciprocal changes in *F_LC_*, the position of the telomeres were not significantly different from the experimental data, with 75.2% and 62.3% of analysed time points at 25°C, and 69.1% and 68.7% at 37°C in the periphery, for Tel3R and Tel14L, respectively (Fig. 4B(d,e), see Fig. S8B and Table S8 for other force changes). Hence, the heteropolymer model was able to simulate the experimentally determined telomeric positions with much greater accuracy than the uniform model.

In both types of analyses, heteropolymers with *F_LC_* changes of ≥5x resulted in an overall compaction of the genome. This is visible from the median 3D distances, which are reduced in comparison to the uniform simulation (Fig. 4A(f,g) vs. Fig. 4A(e)), and from the more central location of the telomeres (Fig. 4B(d,e) vs. Fig. 4B(c)). In both analyses, a reciprocal factor of 5 resulted in the best match to experimental data (Fig. 4A,B, Fig. S8A,B, Table S7,S8). Thus, all subsequently shown simulations used the combinations of 5x / 0.2x to represent 25°C, or 0.2x / 5x to represent 37°C, as *F_LC_* scaling factors for S3 and S4 segments, respectively.

These results demonstrate that the fit between model and experimental data is markedly improved when simulating the chromatin fibre as a heteropolymer with differential *F_LC_*. Heterogeneous mobility of chromatin segments is thus a plausible mechanism shaping chromosome conformation in yeast nuclei.

## DNA-DNA contacts captured by Hi-C and simulation

To gain detailed genome-wide insights into chromosome conformation, we performed Hi-C (*7, 67–71*) in duplicate on yeast cultures (1) grown at 25°C, and (2) grown at 25°C and then shifted to 37°C for 15 min. This allowed us to experimentally determine the genome-wide chromatin contacts at the same growth conditions for which the chromatin states had been determined (Fig. S9). To ensure high quality of the data, we carefully optimised the Hi-C protocol for correct ligation junctions (Fig. S9A,B) and analysed the resulting data for the lack of bias between temperature conditions (Fig. S9C) and for reproducibility (Fig. S9D). We digested the DNA with *Hin*dIII, which in *S. cerevisiae* produces fragments of an average length of 2.7 kb.

We mapped the Hi-C sequencing reads to the yeast genome, and filtered out experimental artefacts, PCR duplicates and physically linked segment pairs using the HiCUP pipeline (*72*). Across the four experiments, this resulted in 12 million valid, unique read pairs (Documents S4-S7). We then plotted and analysed the data at the resolution of individual restriction fragments, i.e. without binning (Fig. 5A). The full contact maps bear the hallmarks of those previously published for *S. cerevisiae:* Strong clustering of the centromeres, weaker clustering of telomeres, and only moderate enrichment of intra- versus inter-chromosomal contacts (*7, 73–75*).

**Fig. 5.**
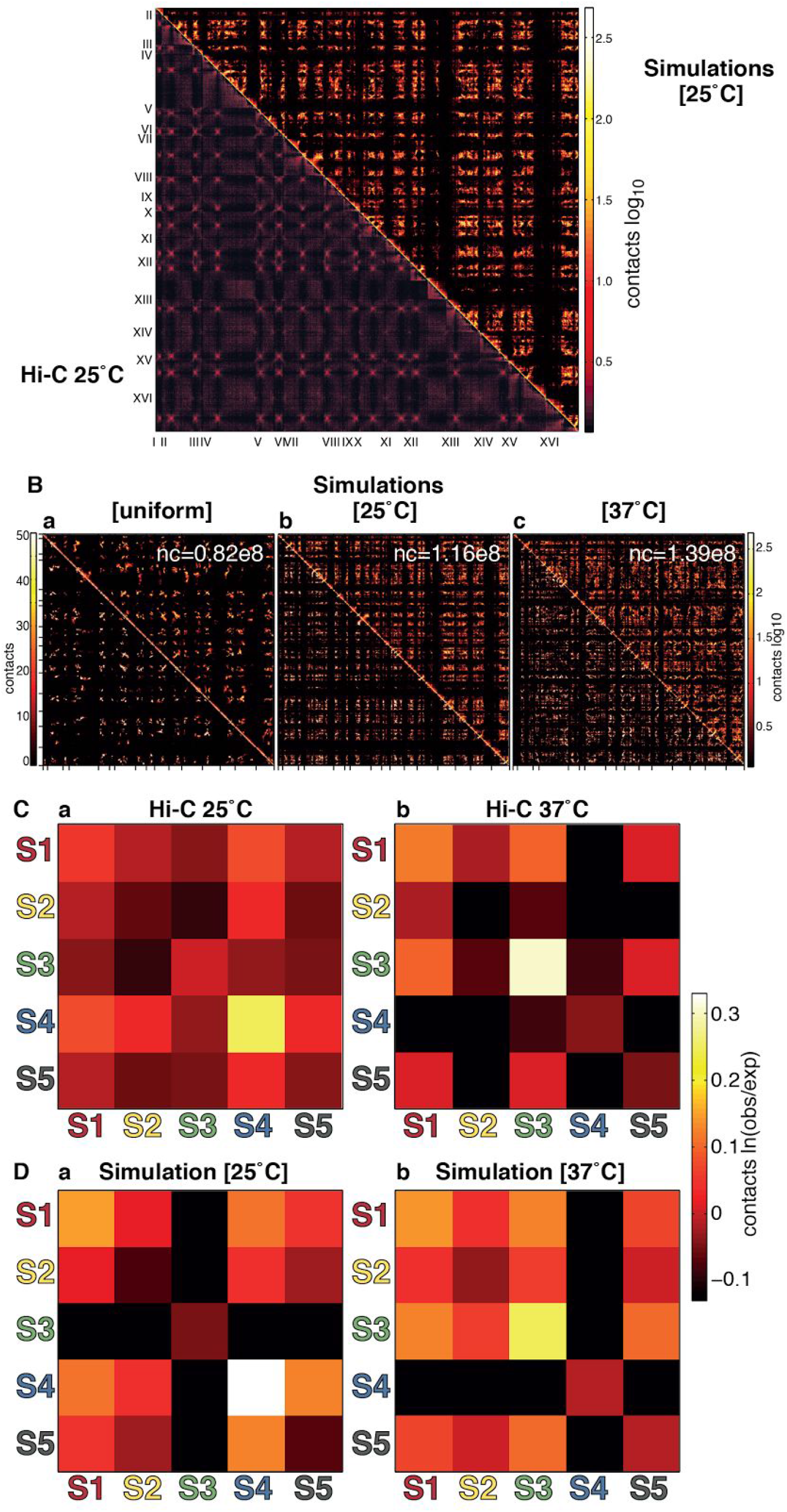
DNA-DNA contact maps of *S. cerevisiae* by Hi-C and simulation. **(A)** Full contact maps of Hi-C at 25°C (below diagonal) and simulations representing [25°C], sampled every 100 time steps (above diagonal). Both are drawn with single-fragment resolution. **(B)** Full contact maps of simulations representing different setups. nc, number of all contacts recorded in 30,000 time points. **(C)** State-wise contact maps of the Hi-C data, from both growth temperatures (25°C and 37°C), normalised by the number of expected contacts as calculated from the proportion of segments in each state. **(D)** State-wise, normalised contact maps of the simulation data.

We analysed the simulations in an analogous manner, recording the incidence and position of contacts between segments throughout the simulation (see SI for details). Each of the three simulations gave rise to distinct patterns of contacts (Fig. 5B). The uniform model (Fig. 5B(a)) gave rise to markedly fewer contacts in the same number of time steps, confirming our observation that heterogeneous mobility achieves greater compaction of chromosomes (Fig. 4).

To understand the influence of protein occupancy on genome organisation, we asked whether chromatin contacts are equally distributed across all chromatin states. After developing a method to map chromatin states to *Hin*dIII fragments (Fig. S4B and SI text), we calculated state-wise contact maps for each growth and simulation condition (Fig. 5C,D and SI). The state-wise contact maps of the experimental and simulation data are remarkably similar in a number of important aspects. Of all state-wise contacts, the highest contact frequencies at 25°C are between *Hin*dIII fragments (Fig. 5C(a)) or model segments (Fig. 5D(a)) of the S4 state. At 37°C, the highest contact frequencies in each case are between fragments or segments in the S3 state (Fig. 5C(b), Fig. 5D(b)). At both temperatures and in both simulations, the highest number of contacts are intra-state in the state with the highest protein occupancy (as determined experimentally) and the lowest mobility (implemented as the lowest *F_LC_* in the model). At the same time, the segments with low protein occupancy and high mobility (S3 at 25°C, S4 at 37°C) have a moderate intra-state contact frequency but show clearly reduced contacts to all other chromatin states.

Interestingly, it is not the fast-moving segments that interact most frequently (Fig. 5D). Instead, it is the slow moving segments that have the highest number of contact, and our simulations indicate that this is a result of close spatial proximity (Fig. 6). In addition, the state-wise contact maps demonstrate that there is a clear spatial separation of the states with high protein occupancy and low mobility from those with low protein occupancy and high mobility, indicated by the ‘black cross’ of low contact frequencies, most clearly seen in Fig. 5C(b) and 5D(a,b).

**Fig. 6.**
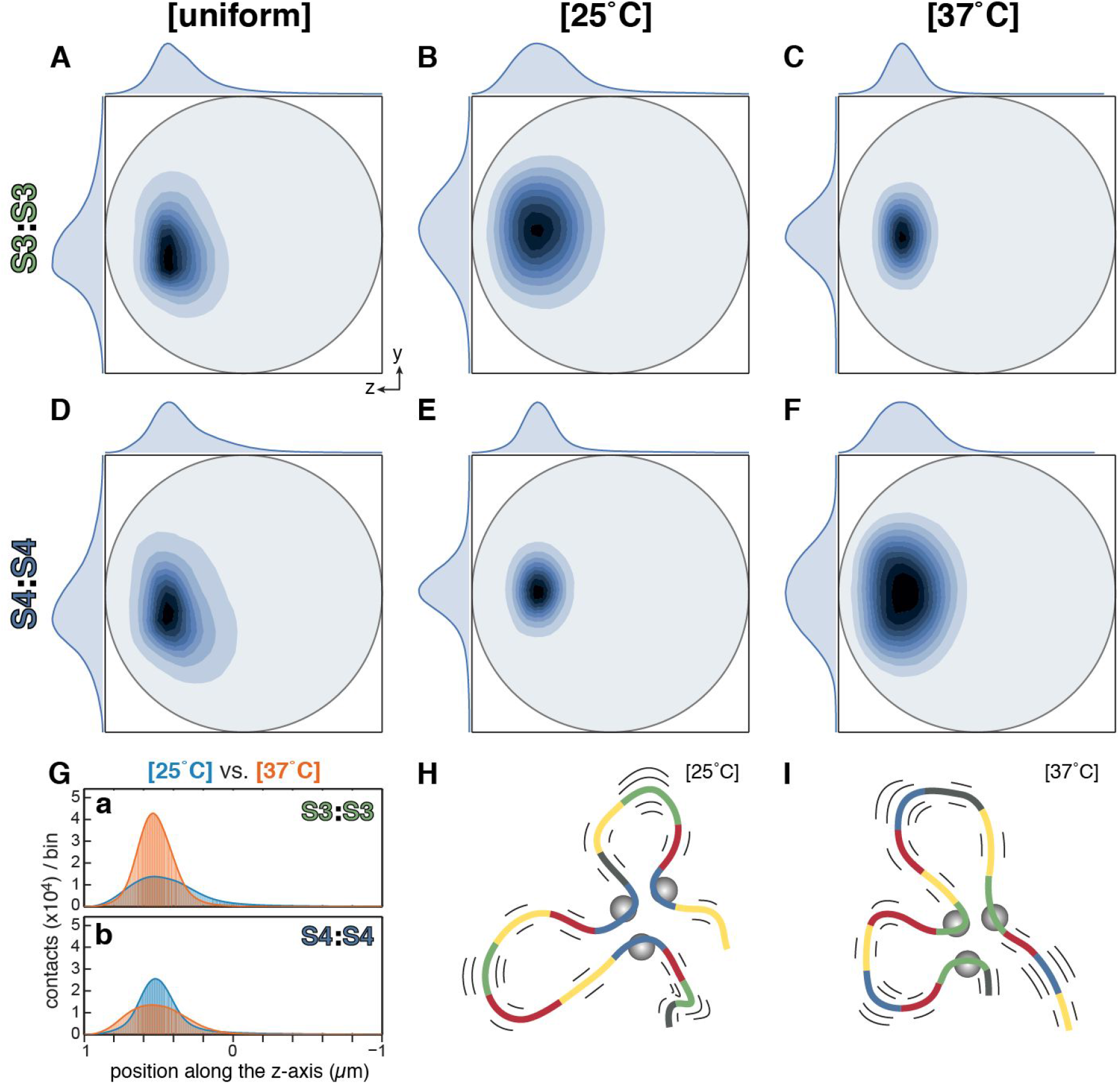
Positions of intra-state contacts in the simulations. **(A-F)** 2D projection density plots of spherical yeast nuclei show the positions of simulated contacts between segments of the same state, in 3 x 1 million time steps (not sampled). Nuclei are oriented as in Fig. 3A. The distributions along the projected axes are given outside the box. The plots are normalised to the highest density in each panel. (A-C) S3 intra-state contacts; (D-F) S4 intra-state contacts. (A,D) uniform model; (B,E) with [25°C] conditions; (C,F) with [37°C] conditions. **(G)** Quantitative histograms of the contacts along the z dimension, in 500 bins of 4 nm width. Profiles at [25°C] (cyan) and [37°C] (orange) are overlaid. They demonstrate the strong clustering of S3:S3 contacts at 37°C **(a)** and of S4:S4 contacts at 25°C **(b). (H,I)** Schematic of how differences in mobility caused by increased binding of poised proteins at S4 segments at 25°C (H), or S3 segments at 37°C (I), leads to clustering in space. Grey spheres represent poised protein complexes; the number and length of the flanking black lines symbolise the degree of segment mobility.

We found that the patterns of the state-wise contact maps are very robust with regards to simulation setup. Different arrangements of the chromosomes around the spindle pole body, sizes of the rDNA, sampling and stochastic repeats produce a different total number of contacts, but the patterns of the normalised state-wise contact maps are virtually indistinguishable, as long as the same statewise *F_LC_* factors are applied (Fig. S6C-D and not shown).

To determine how well the simulated contacts matched the experimental results, we calculated Pearson correlation coefficients between the state-wise contact maps (Table 1): Hi-C data derived from different temperature conditions show no significant correlation, indicative of the significant genome rearrangements that occur after heat shock. Similarly, the [25°C] and [37°C] simulations show no correlation. However, there is high correlation between the Hi-C data and the simulation data at equivalent conditions, indicating that variations in chromatin mobility we implemented in the model shape genome organisation. The heat shock response is likely to induce changes in mobility due to the observed concerted redistribution of protein binding. By this measure, the overall protein occupancy of a chromatin segment alters its mobility and thereby is a significant determinant of 3D genome organisation in yeast.

**Table 1.**
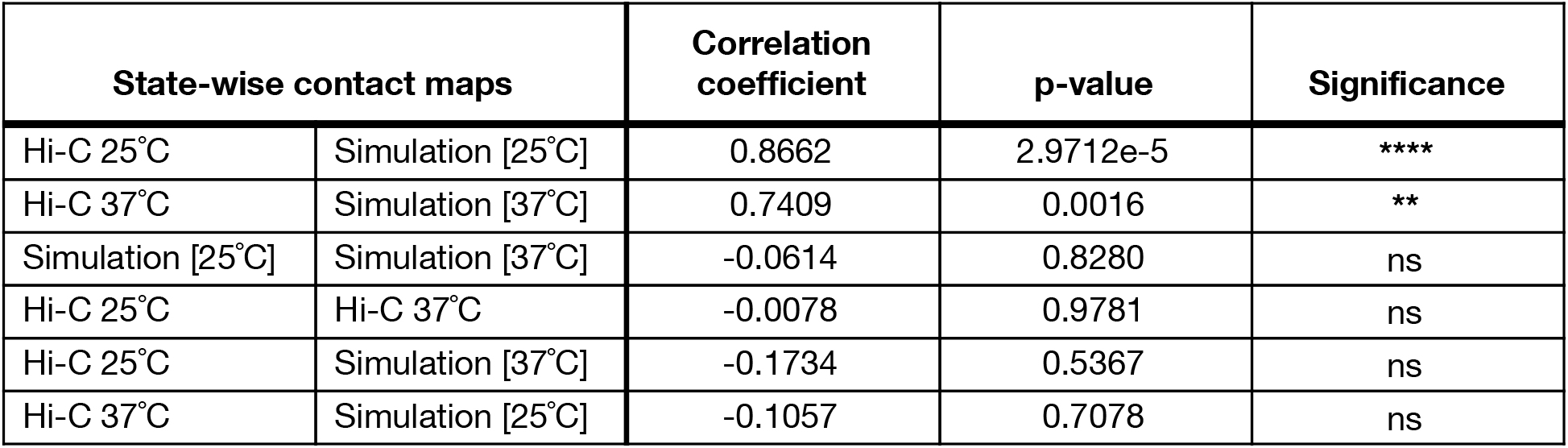
Correlation of state-wise contact maps. Pearson correlation coefficients calculated in MATLAB between two vectors of size 15 (the numerical values of the upper-right triangular parts of matrices depicted in Fig. 5C,D). The rows are sorted by correlation coefficient; ns, not significant.

## Poised genes cluster in 3D

To determine where in the nucleus the simulated contacts occurred, we visualised the position of the intra-state-contacts in 2D projections (Fig. 6, Fig. S10). In all cases, the majority of the contacts are restricted to one half of the nucleus, due to the tethering of the centromeres to the spindle pole body, with the nucleolus occupying the opposite side (*58*). We quantified the level of clustering for all intra-state contacts (see Table S9, Table S10 and SI for details). In the uniform simulations, contacts are the most disperse, confirming the lack of compaction in the homopolymer (as in Fig. 4, Fig. 5B). In addition to this compaction, which affects the chromosomes in their entirety, we observe state-specific clustering of genes in 3D, where genes of one state, dispersed between all 16 chromosomes, preferentially co-localise. Between [25°C] and [37°C], antagonistic clustering occurs: Intra-S3 contacts cluster strongly at [37°C], and intra-S4 contacts cluster strongly at [25°C] (Fig. 6C,E,G), with a 63- and 43-fold change respectively (Table S10, columns 5&6). For comparison, contacts in the entire genome change by only ~1.5-fold (Table S10, last three rows). Preferential clustering is visualised in Fig. 6H,I: In both simulated temperature conditions, the slow-moving, poised segments come together in transient clusters, located at a distance from the nuclear envelope, towards the centre of the nucleus.

## Relocalisation of activated genes towards the nuclear periphery

In yeast, several genes have been described to relocate to the nuclear periphery upon activation and to interact with nuclear pore complexes, aiding the export of mRNA into the cytoplasm (*76–81*). An example is the gene coding for the stress-induced disaggregase HSP104 (*82, 83*), which moves to the nuclear periphery upon induction (*84*). Several molecular mechanisms have been proposed (*53, 76, 80, 84–86*), but the phenomenon is not yet fully understood (*87*). Seeing that our models resulted in changes in location of S3 and S4 genes, we set out to test whether the changes in physical properties of the heteropolymer would suffice to deliver individual genes to the periphery.

We determined the simulated positions of the segment corresponding to the *HSP104* gene at both temperatures and plotted the distribution of positions in radial density plots (Fig. 7A). These density plots show that, at 25°C, the gene is located within a small area and at distance from the nuclear periphery (Fig. 7A, [25°C]). At 37°C, the position of the gene has changed, showing a broader distribution, closer to the periphery (Fig. 7A, [37°C]). The number of times the gene was found within the peripheral area (outside of the dashed circle) more than doubled (Fig. 7C,F) (p<2.2e-16, Wilcoxon rank sum test). Confocal microscopy imaging had shown a very similar increase in peripheral location of the *HSP104* gene upon induction (*84*) (Fig. 7E).

**Fig. 7.**
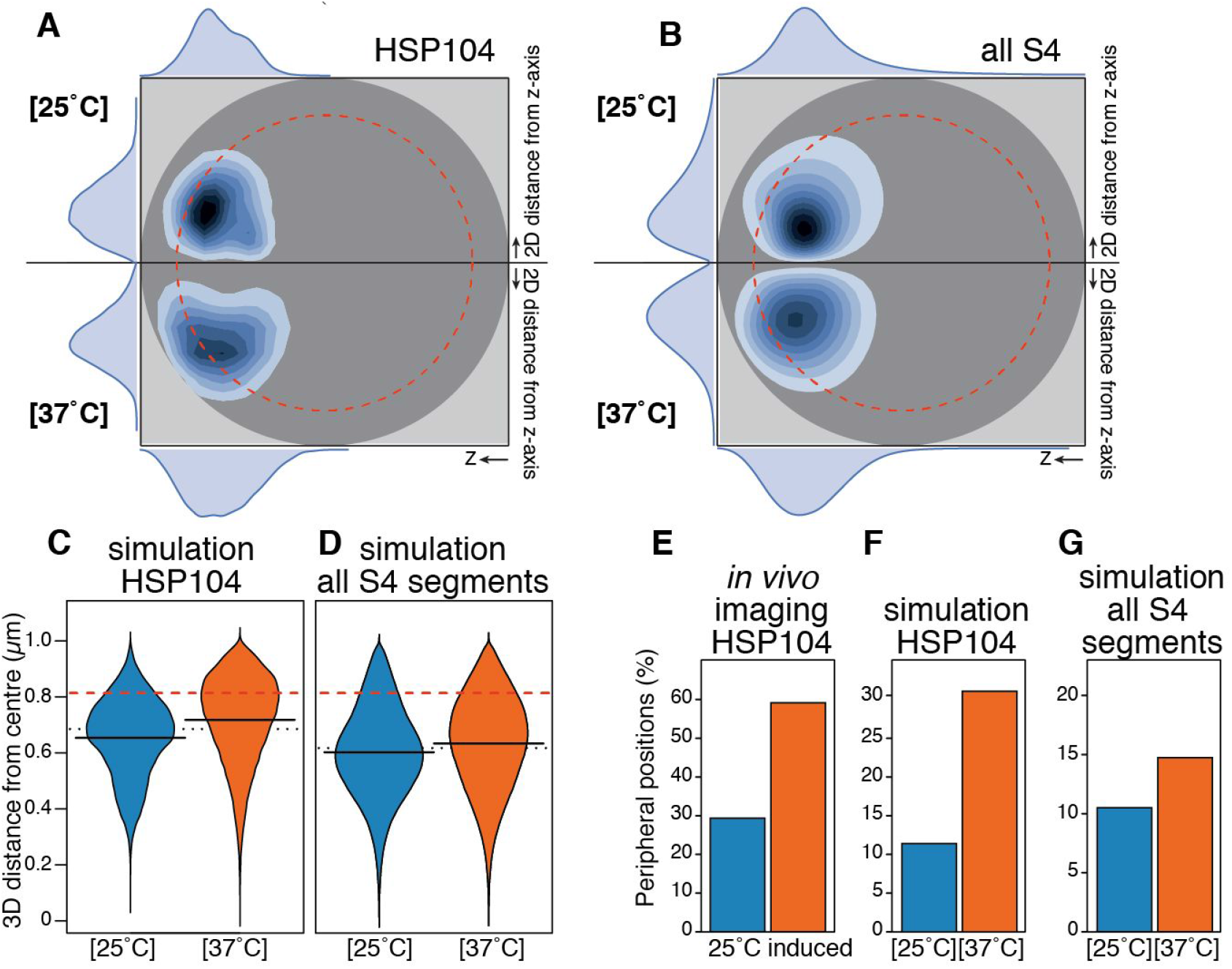
Peripherality of heat-shock gene *HSP104* and all genes in S4 before and after induction. **(A, B)** Plots of the radial density (probability map) around the z-axis of the simulated positions of the segment representing the *HSP104* gene, which belongs to S4, at 25°C (top) and at 37°C (bottom). The SPB is located at the center to the left. The peripheral zone, defined as covering the outermost 33% of the nuclear area, is indicated (dashed red circle). **(B)** The location of all S4 gene segments, plotted as in (A).**(C,D)** Distribution of the linear 3D distances of *HSP104* or all S4 segments from the nuclear centre. **(E)** Corresponding experimental values of *HSP104* (modified from (*84*)) of non-induced and induced cells. **(F,G)** Simulation data, quantified from (A,B).

This difference in location between the two temperature conditions can be seen in the collective shift of distribution of all S4 genes (Fig. 7B,D,G). The difference in peripheral location is highly significant (Fig. 7D,G; p<2.2e-16). Simulating chromosomes as heteropolymers with differential and changing mobility thus allows us to propose this as a general mechanism facilitating the relocation of activated genes to the periphery without the requirement for additional energy sources or specific factors.

## Discussion

A growing body of evidence has shown that genome architecture is closely linked to gene expression and, in higher organisms, cellular differentiation (*11, 88–90*). We now propose a novel mechanism which leads to chromatin states showing distinct spatial organisation. Here, the organisation results from the heterogeneous dynamic behaviour of the chromatin fibre, which is based on the heterogeneous protein occupancy. Of note, it is not the activity of a small number of specific proteins that shape genome organisation, but the cumulative effect of all chromatin-associated proteins that affect the mobility of each genome segment.

The definition of chromatin states allowed us to determine that a large proportion of chromatin-associated proteins, not just RNA polymerase subunits, are poised. This poising is visible at both growth temperatures, and involves rapid and coordinated movement of 42% of chromatin-associated proteins during the transition. At each temperature, the poised proteins are bound to the genes which are currently less active, but which will need to be transcribed quickly if conditions change. It is easy to envision that this widespread poising greatly increases the cell’s ability to quickly activate gene expression, something that is especially of benefit to a free-living, single celled organism encountering unbuffered and rapidly changing external conditions.

The heterogeneous mobility that we infer has several important consequences. Firstly, it leads to an overall compaction of the genome, as seen by the position of the telomeres and 3D distances of chromosomal loci (Fig. 4) and the clustering of contacts (Table S9-S10: in the ‘all contacts’ rows, the uniform conformation is always less compact than the two heteropolymeric ones). This is in addition to the compaction of the chromatin fibre achieved by nucleosomes. The general compaction we observe keeps most of the genome at a distance from the nuclear envelope, leaving more of the peripheral region free for induced genes.

Secondly, our simulations reveal that changes in the distribution of segment mobilities lead to re-organisation of the genome. Specifically, they show that at 37°C, the S4 genes are located closer to the periphery. This represents an astonishingly simple mechanism to re-locate active genes to the nuclear periphery. We are aware that other specific interactions are also involved (e.g. between transcription factors or mRNA binding proteins and the nuclear pore complex (NPC) (*85, 91*)). Interestingly, there are exceptions: the relocation of osmotic shock response genes to the nuclear periphery has been shown to occur even in the absence of nuclear pore complex proteins (*78*). This supports the contention that our mobility-based mechanism can act independently of known specific mechanisms. We envisage that induced genes are primarily delivered to the periphery by our mechanism, and retained there through interactions with the NPC. This could explain the quantitative differences between the *in vivo* data and our simulation outcomes (Fig. 7 E,F).

Finally, the spatial clustering of the poised state segments is relevant in the context of the observed clustering of genes and proteins that lead to the formation of transcription factories (*78*). While still speculative, this clustering offers the opportunity to help all proteins that are constituents of transcription factories to find each other prior to activation of transcription.

Our results demonstrate that, given the known architectural constraints of the *S. cerevisiae* genome, the differential mobility of chromatin segments is a significant determinant of overall 3D genome organisation. This requires a highly dynamic system, in which chromosomes are in constant motion. The self-organisation we observe does not rely on protein-protein interactions, such as those mediated by cohesin (*92*) or CTCF in higher eukaryotes (*93*). Intriguingly, it has recently been shown that the major structural compartments in the genomes of mouse embryonic stem cells remain intact even when CTCF is depleted (*94*), (*95*), indicating that other, CTCF-independent, mechanisms are involved (*96*). These mechanisms remain to be elucidated and could include self-organisation by differential mobility.

The mechanism we propose is akin to granular convection, also known as the Brazil nut effect, in which objects of different size and similar density separate spatially when shaken (*97, 98*). In addition to phase separation caused by size, shape and mass, particles that have different diffusion properties can also separate, as has recently been demonstrated theoretically and using simulations for monomers (*99*), polymers (*100–102*).

As mentioned when introducing the model, we chose the compound Langevin Force as an intentionally non-specific means of changing the mobility. The precise components that influence chromatin mobility are the subject of active ongoing studies. Several aspects of protein physics, chemistry and biology are of importance, and it is the joint effect of all these influences that determines the heterogeneity of chromatin segment diffusion. High protein occupancy, as in the poised segments we identified, directly increases the mass and diameter of the chromatin segment. It furthermore increases the opportunities for transient interactions of the chromatin segment with neighbouring structures, thus reducing their effective diffusion speed (*103–105*).

Effective diffusion can be affected by the charge, the secondary and tertiary structure of bound proteins. Extended, highly unstructured protein domains could result in particles with a higher viscosity (*106, 107*), and thus reduce mobility. We calculated the internal disorder score of all chromatin associated proteins and compared how these scores varied across the chromatin states, and found a significant enrichment of disordered domains in the poised state at both temperatures, corroborating our model that poised segments move more slowly (Sewitz and Lipkow, *unpublished*). Recently, intrinsically disordered domains of the *S. cerevisiae* transcription factor Mig1p have been reported to stabilise proteins clusters by entropic depletion (*108, 109*), forming transient bridges between DNA strands, and reducing diffusion of the protein-DNA complexes (*108, 109*). Interestingly, Mig1p is repressive, thus its binding simultaneously reduces gene expression and mobility. This provides experimental support for our postulate that poised genes, which are currently not highly expressed, have a reduced mobility.

Even enzyme activity can affect the undirected, stochastic mobility of chromatin segments: It has recently been proposed that the heat produced by ATP hydrolysis as a side effect of transcription and chromatin remodelling can lead to increased stochastic fluctuations. In simulations of 1 Mb resolution, this achieves the separation of gene-dense (active) from gene-poor (inactive) chromosomes (*100, 110, 111*), albeit at unphysiogically high temperature differences (20-fold). Interestingly, in finer models of long homopolymers, this temperature difference can be reduced to more realistic values (*100, 101*). The recent report that *Drosophila* topological domains separate by transcriptional state (*96*) lends further support to this idea. All this supports the concept that numerous factors are involved in modifying chromatin mobility.

Lastly, several recent reports highlight how the physical process of phase-separation contributes to chromatin organisation (*112, 113*). Looking forward, integrating this with the role played by noncoding eRNAs and the mechanism describing how multivalency regulates RNA granule size (*114, 115*), promises to expose the complexity of biophysical mechanisms that together work to carefully control genome organisation.

## Acknowledgments

We are indebted to Jonathan Baxter, Stephanie Schalbetter, Job Dekker and Jon-Matthew Belton for sharing their unpublished yeast Hi-C protocol and advice. We thank Kristina Tabbada for sequencing, Charlotte Kaplan Galimow and Matthias Benoit for help with generating yeast strains, and Jonathan Houseley for advice. We thank Christophe Zimmer for sharing the code of their published yeast model, Pascal Carrivain for discussions on Langevin equations and ODE, Jonathan Cairns and Anne Segonds-Pichon for advice on statistics, and Guillaume Filion for providing the HMMt package and advice. The Hi-C data is archived at the Gene Expression Omnibus (http://www.ncbi.nlm.nih.gov/geo/), accession number XXX.

We acknowledge funding by a Royal Society University Research Fellowship UF071175 & UF120022 (KL), Royal Society Research Grant RG080501 (KL), Microsoft Research Faculty Fellowship contract 2009-021 (KL), Apple Research & Technology Support (KL), the Babraham Institute (KL, SAS), Islamic Development Bank (ZF), King Saud University (LA), CNPq (OJBB), EU FP7 contract 201142 (SGO), Biotechnology and Biological Sciences Research Council UK grant BBS/E/B/000C0405 (PF), MRC MR/L007150/1 (PF), ERC Advanced Grant DEVOCHROMO (PF), Agence Nationale pour la Recherche ANR *ANDY* (KB), Institut Universitaire de France (KB).

## Supplementary Materials

Materials and Methods

Figures S1-S10

Tables S1-S9

Movie S1

Documents S1-S7

External Database S1

Author Contributions

References 116-126

